# Recapitulation of Human Pathophysiology and Identification of Forensic Biomarkers in a Translational Swine Model of Chlorine Inhalation Injury

**DOI:** 10.1101/2022.02.09.479576

**Authors:** Satyanarayana Achanta, Michael A. Gentile, Carolyn J. Albert, Kevin A. Schulte, Brooke G. Pantazides, Brian S. Crow, Jennifer Quiñones-González, Jonas W. Perez, David A. Ford, Rakesh P. Patel, Thomas A. Blake, Michael D. Gunn, Sven E. Jordt

## Abstract

**Rationale:** Chlorine gas (Cl_2_) has been repeatedly used as a chemical weapon, first in World War I and most recently in Syria. Life-threatening Cl_2_ exposures frequently occur in domestic and occupational environments, and in transportation accidents. There is a knowledge gap in large animal models of Cl_2_-induced acute lung injury (ALI) required to accurately model human etiology and for the development of targeted countermeasures

**Objective:** To develop a translational model of Cl_2_-induced ALI in swine to study toxicopathophysiology and identify biomarkers useful for forensic analysis.

**Methods:** Specific pathogen-free Yorkshire swine (30-40 kg) of either sex were exposed to Cl_2_ gas (≤ 240 ppm for 1 h) or filtered air under anesthesia and controlled mechanical ventilation.

**Results:** Exposure to Cl_2_ resulted in severe hypoxia and hypoxemia, increased airway resistance and peak inspiratory pressure, and decreased dynamic lung compliance. Chlorine exposure resulted in increased total BALF and neutrophil counts, vascular leakage, and edema compared to the control group. The model recapitulated all three key histopathological features of human ALI, such as neutrophilic alveolitis, deposition of hyaline membranes, and formation of microthrombi. Free and lipid-bound 2-chlorofatty acids and chlorotyrosine-modified proteins (3-chloro-L-tyrosine and 3,5-dichloro-L-tyrosine) were detected in plasma and lung after Cl_2_-exposure.

**Conclusions:** The translational model developed in this study replicates key features of humans exposed to Cl_2_ and is suitable to test medical countermeasures. Specific biomarkers of Cl_2_ exposure have been identified in plasma and lung tissue samples.

**Take home message:** We developed a swine model of chlorine gas-induced acute lung injury that exhibits several features of human acute respiratory distress syndrome. We validated chlorinated fatty acids and protein adducts in plasma and lung samples as forensic biomarkers.

## INTRODUCTION

Chlorine gas (Cl_2_) is a highly reactive toxic halogen. Chlorine gas is safe for several domestic, occupational, and industrial purposes when used at appropriate concentrations. However, it can be toxic when used inappropriately. Cl_2_ has been used as a chemical weapon since World War I and most recently in chlorine bomb attacks in Syria [1, 2]. The UN-supported OPCW fact-finding missions proved that Syria used Cl_2_ allegedly on the civilian population [2]. Since Cl_2_ can be easily synthesized from simple ingredients, Cl_2_ has resurged as a potential chemical warfare agent. Unfortunately, biomarkers that can confirm Cl_2_ exposure have only been validated in *in vitro* and rodent models [3-6].

Train derailments have been sources of accidental exposure to toxic concentrations of Cl_2_. Several major cargo derailments and other transportation accidents have occurred throughout the world, resulting in morbidity and mortality [7-11]. It is estimated that chlorine accounts for 84% of the total toxic inhalation hazards (TIH) transported every year [12]. The train derailment incident in Graniteville, SC in 2005 is considered one of the largest Cl_2_ accidents in the United States. In that incident, 9 people died and 200 were admitted for inhalational pulmonary injuries [7, 9, 10, 13]. In addition to acute casualties, long-term inflammation and hyperreactivity have also been reported in victims of the Graniteville derailment [10, 14, 15].

The pathophysiology and symptoms of acute lung injury (ALI) caused by Cl_2_ are highly variable, and based on concentration and exposure time [16-19]. The common acute symptoms of Cl_2_ inhalation are consistent with an acute respiratory impairment, and include running nose, sore throat, cough, choking, tightness of chest, labored breathing, bronchospasms, and wheezing [7, 17, 20]. Exposure to higher concentrations of Cl_2_ for a prolonged period may result in non-cardiogenic edema. At concentrations above 1000 ppm, Cl_2_ can be fatal to humans within a few minutes [18, 21].

Due to increased concerns about the use of Cl_2_ as a potential chemical threat agent and the likelihood of accidental and occupational exposures, there is an immediate need for therapeutic agent(s) against Cl_2_-induced ALI. Plume models have predicted the gravity of a Cl_2_ attack or accidental release [22-24]. Despite the use of Cl_2_ as a chemical warfare agent since World War I, there is no specific antidote for this powerful choking agent except symptomatic treatment [17, 19]. Although several potential therapeutic agents have been tested for efficacy in Cl_2_-induced ALI in different preclinical animal models, none have been approved [19, 25, 26]. Due to ethical and feasibility issues surrounding clinical trials of chemical threat agents in humans, the United States Food and Drug Administration (FDA) made provisions to approve therapeutic agents under animal rule (21 CFR 314.600) [27]. Animal rule recommends demonstration of therapeutic efficacy in at least 2 animal models, including a higher mammalian species close to humans in phylogeny. Rodent models have been routinely used to screen for the initial therapeutic efficacy of candidate drugs [19, 28]. However, no ideal large animal model that mimics several features of human Cl_2_-induced ALI/ARDS (acute respiratory distress syndrome) has yet been developed, which remains a barrier to testing potential therapeutic candidates. Therefore, *we developed a pig translational model of human chlorine gas-induced ALI, and validated chlorinated fatty acids and chlorinated tyrosine adducts as potential biomarkers of Cl*_*2*_ *exposure*. We chose the pig as a translational model to study chemically induced ALI in humans due to their anatomic and physiologic similarities [29, 30].

## MATERIALS AND METHODS

### Please see additional details on materials and methods in the online supplementary information

#### Preparation of animals, Cl_2_ exposure, and data collection

Specific pathogen-free (SPF) male and female Yorkshire pigs (n =14) (castrated males and intact females) (30-40 kg) were procured, and given at least 48 hours to acclimate. The Duke University Institutional Animal Care and Use Committee (IACUC) approved the procedures in this study.

Animals were anesthetized and mechanically ventilated. After placement of surgical instrumentation and acquisition of baseline data (respiratory and cardiovascular physiologic parameters, and arterial blood gas analysis), animals were randomized to Cl_2_ or filtered room air exposure (n=6/air group; 8/Cl_2_ group). (Two pigs died during or a few hours after Cl_2_ exposure; therefore, data were presented for 6/group.) Cl_2_ was delivered via the inspiratory limb of the mechanical ventilator circuit, and monitored with a chlorine detector (Porta Sense II Gas Leak Detector, AFC International, Inc, DeMotte, IN, USA) at 8-15-minute intervals to ensure delivery of ≤ 240 ppm for 1 hour. Cl_2_ exposure occurred in a negatively pressurized room with adequate ventilation and safety monitoring for leak detection in multiple locations. Figures 1A and 1B show the study paradigm and the schematic of the Cl_2_ exposure system for pigs.

**Figure 1:**
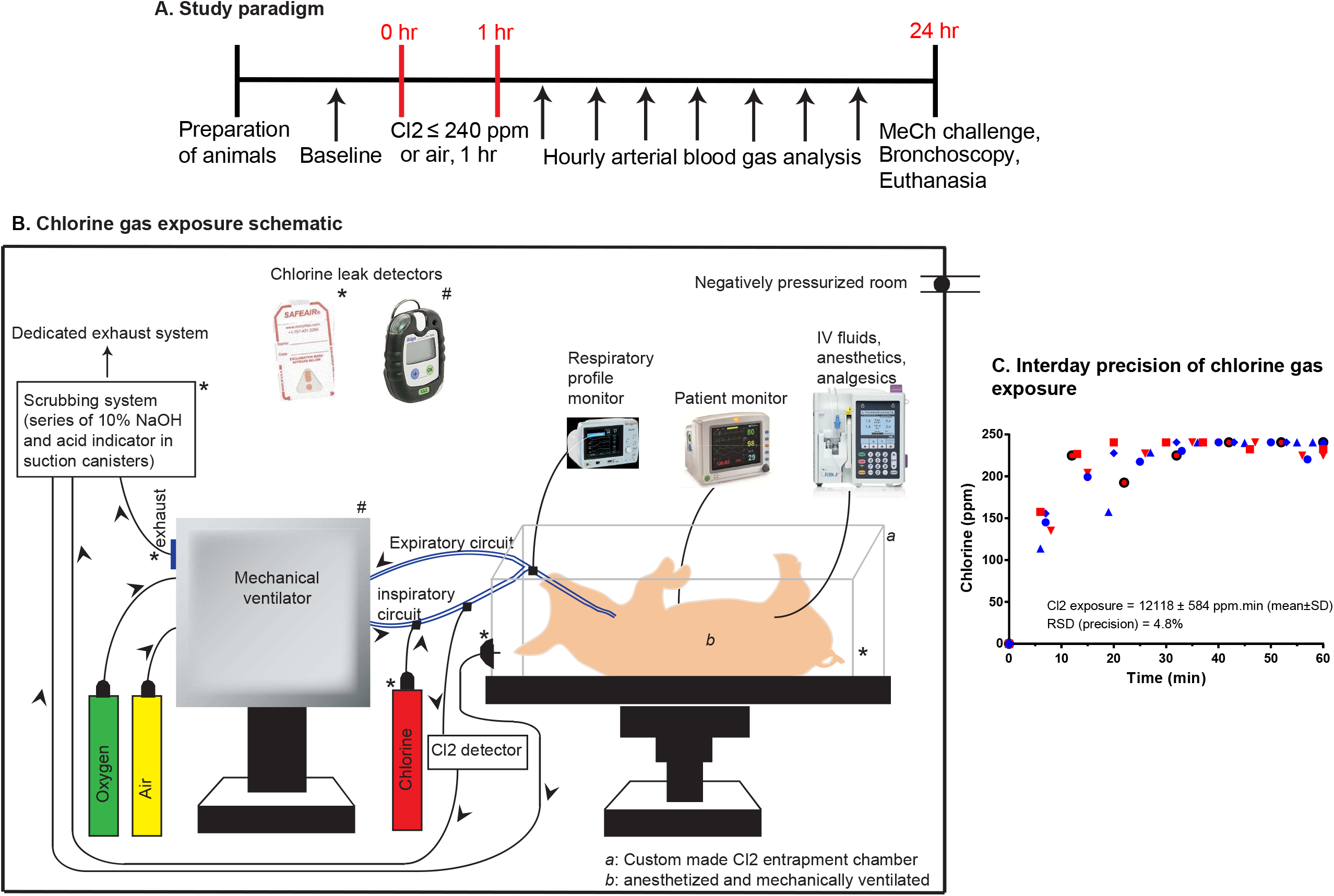
Study design paradigm (1A), chlorine gas exposure schematic (1B), and interday precision of chlorine gas exposure (1C). Anesthetized and mechanically ventilated pigs were exposed to either chlorine gas at ≤ 240 ppm or filtered room air for 1 hour. Chlorine exposure was monitored every 8-15 minutes. Arterial blood gas analysis was performed at hourly intervals, and all key oxygenation and respiratory physiologic parameters were recorded. At 24 hours post chlorine gas or filtered air exposure, methacholine airway challenge was performed, bronchoalveolar lavage fluid was collected with the aid of bronchoscopy, and pigs were then euthanized to conduct gross necropsy and sample collection.

We collected complete blood counts (CBC) and serum chemistry panel values at baseline and at 24 hours post exposure. The key respiratory physiologic parameters were recorded and arterial blood gas analysis was performed at 5 and 30 minutes post exposure, and then subsequently at 1-hour intervals until euthanasia at 24 hours post exposure. Oxygenation parameters, ie, the ratio of the partial pressure of oxygen in arterial blood and fraction of inhaled oxygen (PaO_2_/FiO_2_), oxygenation index, saturation of oxygen in hemoglobin, and arterial-alveolar (A-a) gradient, were primarily considered. Respiratory physiologic parameters, such as compliance dynamics (C_dyn_), airway resistance (Raw), and peak inspiratory pressure (PIP), were assessed at hourly intervals. At 24 hours post Cl_2_ or air exposure, airway mechanics were measured by challenging pigs to increasing concentrations of methacholine hydrochloride (MeCh) (0, 0.5, 1, 2, 4, 8, 16, and 32 mg/mL) (Sigma-Aldrich, St. Louis, MO, USA) via an Aeroneb™ nebulizer (Aerogen Ltd, Ireland). Bronchoscopy was performed to collect bronchoalveolar lavage fluid (BALF) just before euthanasia, and total and differential leukocyte counts were determined as previously published, with some modifications [28].

At the end of the 24-hour period, we euthanized pigs following *AVMA guidelines on euthanasia*, and collected lung tissues for histopathology and diagnostic marker analysis. A board-certified veterinary pathologist performed blinded histopathologic scoring of lung specimens following *an official American Thoracic Society (ATS) Workshop Report:* Features and Measurements of Experimental Acute Lung Injury in Animals, with some modifications (Table S1)[31]. We determined edema using lung wet/dry weight ratio as a surrogate marker. Concentrations of pro-inflammatory cytokine markers were determined in BALF supernatants, serum samples, and lung tissue homogenates using ELISA kits (R&D Systems, Minneapolis, MN, USA).

#### Chlorinated fatty acids (CFAs) in lungs and plasma

Free and total (ie, free + esterified) 2-chlorofatty acids were measured, as previously described, by LC/MS following Dole extraction [32]. Total 2-chlorofatty acids were measured by LC/MS after base hydrolysis, while free 2-chlorofatty acids were not subjected to base hydrolysis before LC/MS analysis. Extractions were performed using 25 µL of plasma spiked with 103.5 fmol of 2-chloro-[d4-7,7,8,8]-palmitic acid (2-[d4]-ClPA) as the internal standard. Lung tissue (20 mg) was pulverized and subjected to Bligh-Dyer lipid extraction [33] in the presence of 103.5 fmol of 2-[d4]-ClPA (internal standard). Half of the lung lipid extract was then analyzed by LC/MS for free 2-chlorofatty acids, and the other half of the extract was subjected to base hydrolysis followed by LC/MS analysis for total 2-chlorofatty acids.

#### Chlorinated tyrosine adducts (CTAs) in plasma and lung tissues

The biomarkers 3-chlorotyrosine (Cl-Tyr) and 3,5-dichlorotyrosine (Cl2-Tyr) were isolated from the pronase digest of plasma and lung tissues samples from air or chlorine exposed pigs by solid-phase extraction (SPE), separated by reversed-phase HPLC and detected by tandem mass spectrometry (MS-MS), following previously published methods [34, 35].

#### Data and statistical analysi

Data are presented as the mean ± standard error of mean estimate (SEM) in figures and as mean ± standard deviation (SD) in tables. Oxygenation and respiratory physiologic parameters are presented as time course scatter plots over 24 hours, and bar graphs represent area under the curve (AUC) from post-Cl_2_ or -air exposure through 24 hours. Unpaired Student’s *t*-test (two-tailed) was performed for comparison of means of 2 groups for statistical analysis unless mentioned (GraphPad Prism version 9 for Windows, GraphPad Software, San Diego, California USA). An α value < 0.05 was considered significant. Statistical significance was denoted as: \**p* ≤ 0.05; \*\**p* ≤ 0.01; \*\*\**p* ≤ 0.001; \*\*\*\**p* < 0.0001.

## RESULTS

We chose a combination of 240 ppm for the concentration of Cl_2_ and 1 hour for time of exposure based on our initial chlorine exposure-response studies. Figure 1C shows interday variability and precision of Cl_2_ exposure. The interday precision of AUC of Cl_2_ exposure was 4.8%.

### Complete blood counts and serum chemistry panels

CBC and serum chemistry panel values are presented in Tables S3 and S4, respectively. Platelets had significantly decreased in both groups at 24 hours post exposure compared to their respective baseline values. Also, the mean platelet count was different between air- and Cl_2_-exposed groups at the 24-hour time point.

Although other parameters had some trends in complete blood counts, they did not reach statistical significance. Among serum chemistry panel values, there was no significant difference between air- and Cl_2_-exposed groups at the 24-hour time point. However, there was a significant difference between values at baseline and the 24-hour time point for glucose, cholesterol and alkaline phosphatase in both air- and Cl_2_-exposed groups. As there are no published references for CBC and serum chemistry in a pig model of Cl_2_ inhalation injury, the data presented here will serve as a reference for future work.

### Acid-base imbalance in Cl_2_ exposed pigs

In the Cl_2_ group, immediately after gas exposure, the pH and HCO3^-^ fell, and partial pressure of carbon dioxide in the arterial blood (PaCO2) increased. Therefore, the initial acid-base imbalance suggests respiratory acidosis with metabolic acidosis. Due to a compensatory increase in HCO3 ^-^, the pH increased, and then decreased toward the end of the study. However, over 24 hour observation, no significant difference in AUCs of pH was noted between the air and chlorine groups; The PaCO_2_ and HCO3 ^-^ were decreased in the chlorine-exposed group whereas lactate values increased (Figure S1). The resulting acid-base imbalance over 24 hour suggests respiratory alkalosis with metabolic acidosis.

### Oxygenation parameters

Pigs in both groups started with identical oxygenation parameters. Cl_2_ exposure resulted in immediate hypoxia and hypoxemia. Pigs in both groups were mechanically ventilated with 21% oxygen, and FiO_2_ was adjusted to maintain at least 80% oxygen saturation in hemoglobin (SpO_2_). Pigs in the chlorine group required higher FiO_2_ values (95% confidence intervals [CI] of AUC, 5.56-6.6) compared to the air group (95% CI of AUC, 5.0-5.1) to maintain SpO2 values above 80% (Fig. 2A). SpO_2_/FiO_2_ values in the chlorine group (95% CI of AUC, 8580-9776) were significantly decreased compared to the air group (95% CI, 10764-10999) (Fig. 2B). PaO_2_/FiO_2_ values in the chlorine group (95% CI of AUC, 5663-6777) were significantly lower than in the air group (95% CI, 9013-9343) (Fig. 2C). The oxygenation index (OI) was significantly increased in the chlorine group (95% CI of AUC, 105-145) vs the air group (95% CI, 49.4-54.3) (Fig. 2D). The arterial-alveolar (A-a) gradient was significantly elevated in the chlorine group (95% CI of AUC, 1303-2020) compared to the air group (95% CI, 224-355) (Fig. 2E).

**Figure 2:**
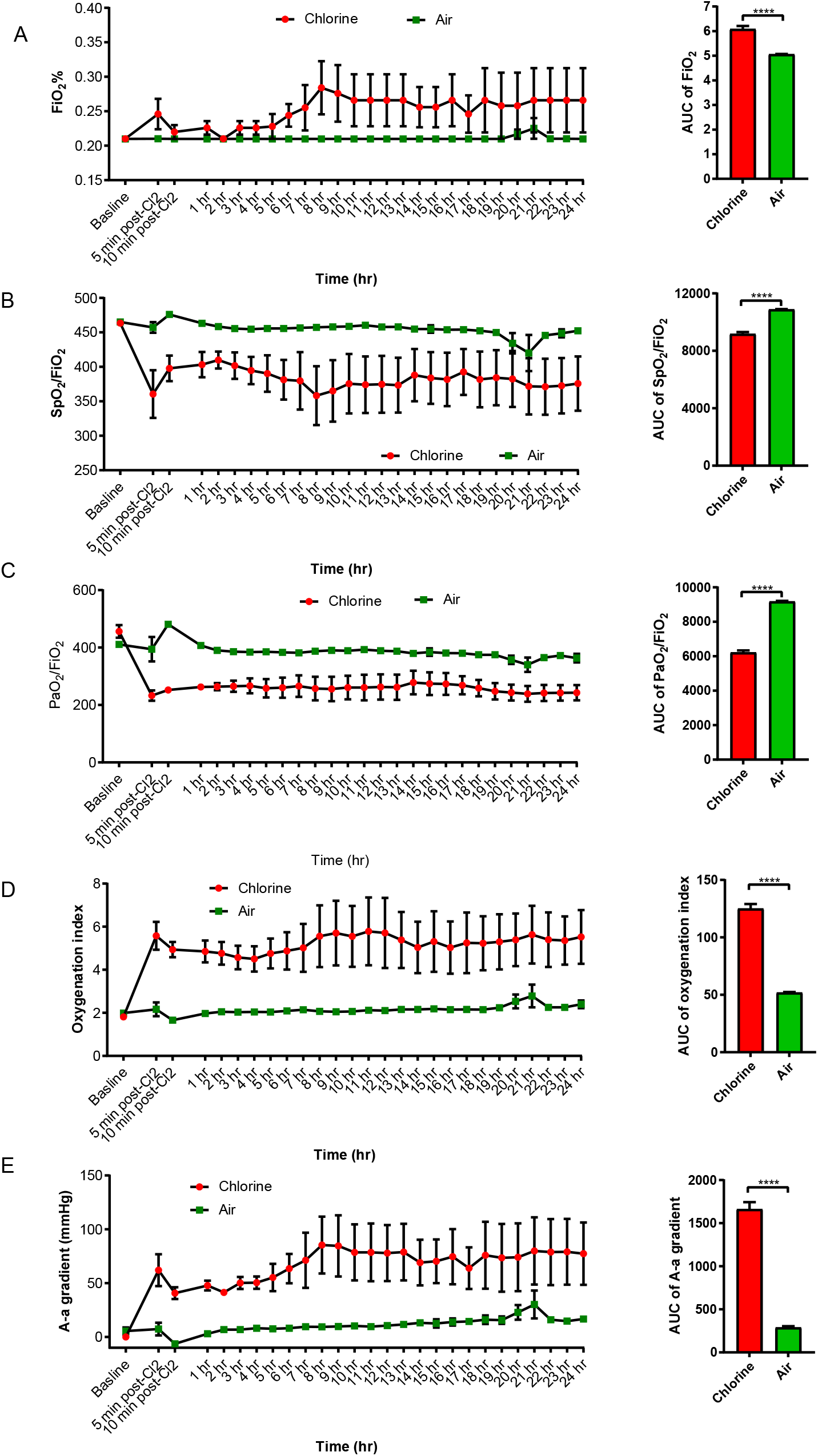
Oxygenation parameters in chlorine-exposed pigs. Anesthetized and mechanically ventilated pigs were exposed to either chlorine gas at ≤ 240 ppm or filtered room air for 1 hour. The fraction of inhaled oxygen (FiO_2_) (Fig 2A), oxygen saturation in hemoglobin normalized by FiO_2_ (SpO_2_/FiO_2_) (Fig 2B), the ratio of partial pressure of oxygen to fraction of inhaled oxygen (PaO_2_/FiO_2_) (Fig 2C), oxygenation index (Fig 2D), and arterial-alveolar (A-a) gradient (Fig 2E) were measured at hourly intervals. The time-course scatter plots show the dynamics of the oxygenation parameters over the 24-hour study period. Bar graphs show the area under the curve over the 24-hour study period, and were compared by unpaired two-tailed *t*-test. ****, *p* < 0.0001.

### Respiratory physiologic parameters

Pigs in both groups had similar baseline respiratory physiologic parameters. Airway resistance was significantly increased in the chlorine group (95% CI of AUC, 332-459) compared to the air group (95% CI, 152-182) (Fig. 3A). Similarly, peak inspiratory pressure was significantly increased in the chlorine group (95% CI of AUC, 645-764) vs the air group (95% CI, 378-419) (Fig. 3B). Compliance dynamics was significantly decreased in the chlorine group (95% CI of AUC, 306-357) compared to the air group (95% CI, 583-673) (Fig. 3C).

**Figure 3:**
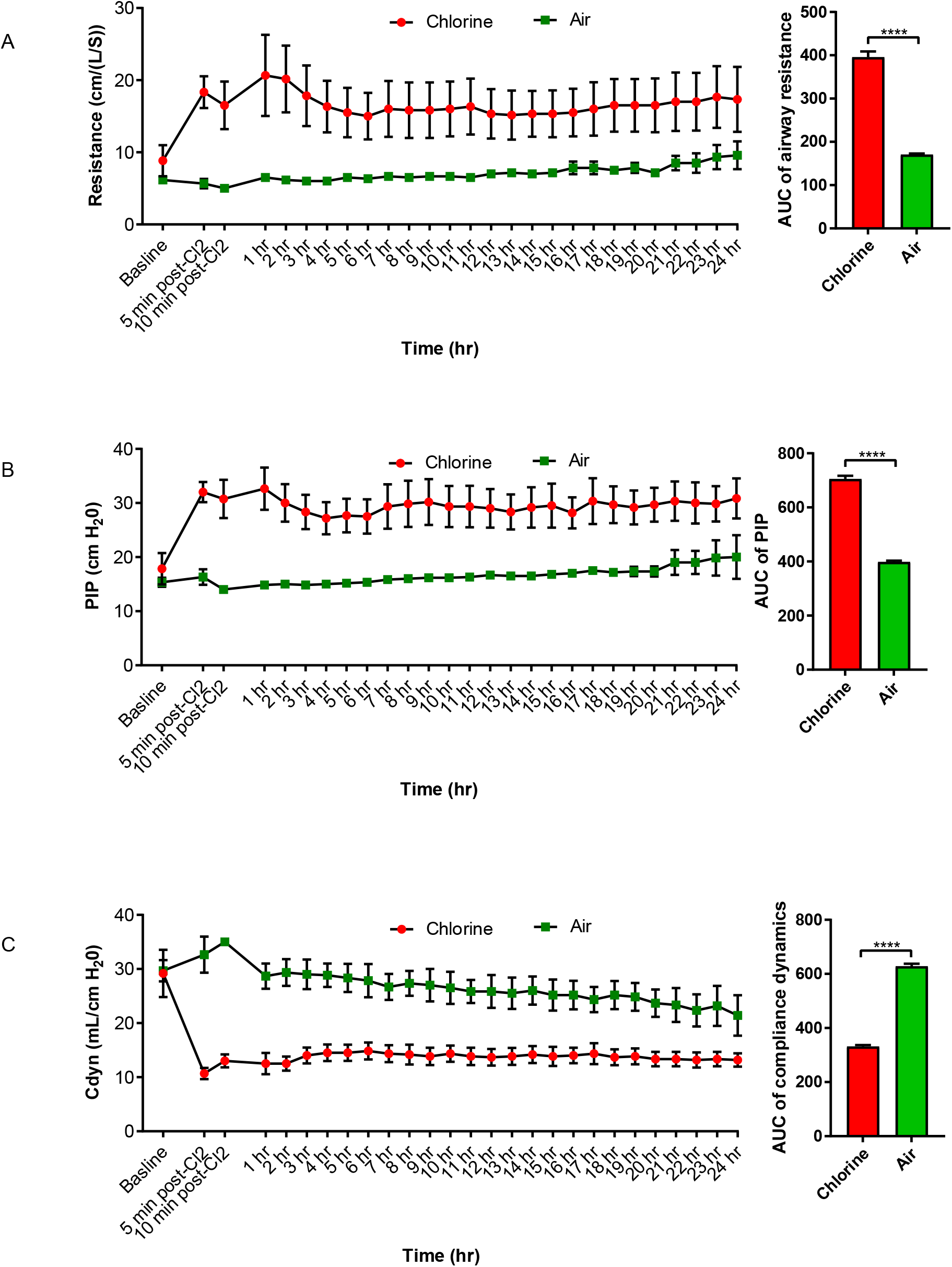
Pulmonary physiologic parameters in chlorine-exposed pigs. Anesthetized and mechanically ventilated pigs were exposed to either chlorine gas at ≤ 240 ppm or filtered room air for 1 hour. Airway resistance (Fig 3A), peak inspiratory pressure (PIP) (Fig 3B), and compliance dynamics (C_dyn_) (Fig 3C) were measured at hourly intervals. The time-course scatter plots show trends in pulmonary physiologic parameters over the 24-hour study period. Bar graphs show the area under the curve over the 24-hour study period, and were compared by unpaired two-tailed *t*-test. ****, *p* < 0.0001.

### Methacholine airway challenge

At 24 hours post exposure, the Cl_2_ group displayed heightened basal pulmonary resistance compared to the air group. Figures 4A-C show responses in airway resistance, PIP, and C_dyn_ with increasing concentrations of methacholine (MeCh). The Cl_2_ group exhibited airway hyperreactivity to inhaled MeCh, whereas the air group tolerated higher concentrations of MeCh.

**Figure 4:**
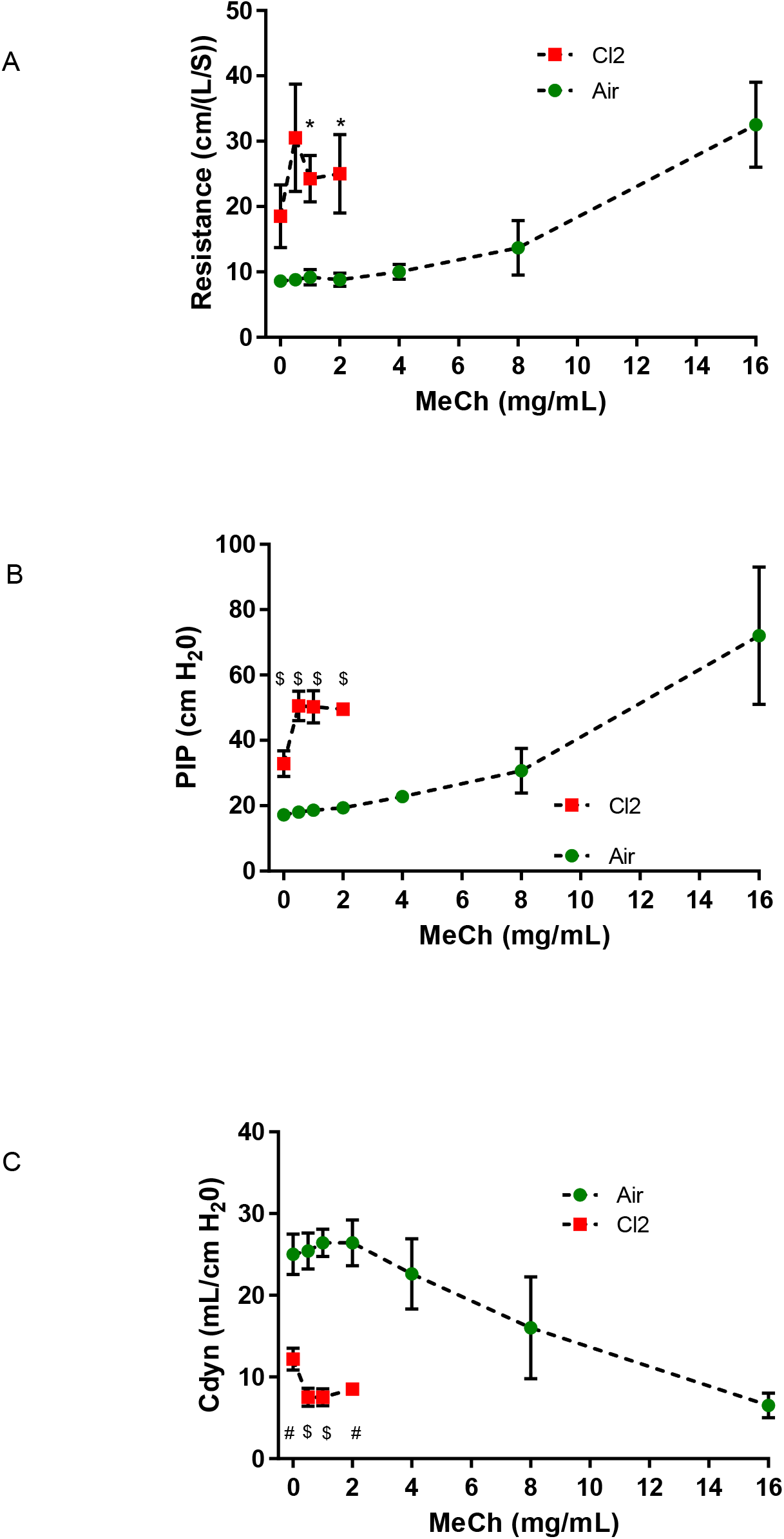
Pulmonary mechanics in chlorine-exposed pigs. Methacholine airway challenge was conducted to assess airway mechanics in chlorine- or air-exposed pigs. Airway resistance (Fig 4A), peak inspiratory pressure (PIP) (Fig 4B), and compliance dynamics (C_dyn_) (Fig 4C) were measured. Unpaired two-tailed *t*-tests were performed between chlorine and air groups at each concentration of methacholine. n=6/Cl_2_-exposed and 5/air-exposed pigs. *, *p* <0.05; ^#^, *p* < 0.01; ^$^, *p* < 0.001.

### BALF total cell counts and differential leukocyte counts, protein leak, and edema

The total BALF leukocyte count was significantly increased in Cl_2_ pigs compared to the air group (Fig. S2A). BALF protein levels and wet/dry lung weight ratio were significantly increased in the Cl_2_ group compared to the air group (Fig. S2B and S2C).

### Pro-inflammatory cytokine markers

IL-6 and VEGF levels were increased in Cl_2_ pigs in all three biological matrices tested, ie, BALF, serum, and lung tissue homogenates (Fig. S2D-F).

### Postmortem examination and histopathologic analysis

Postmortem gross examination of the lungs revealed diffuse lesions of atelectasis and hemorrhage in the Cl_2_ group, whereas the air group had no detectable abnormal findings. However, caudal atelectatic lesions were found in the lungs of both groups. Such lesions are consistent with mechanical ventilator-induced lung injury (VILI). In pigs exposed to air, histopathologic analysis of the lungs revealed normal open alveolar architecture of the parenchyma without any significant lesions. In pigs exposed to Cl_2_, however, lung sections showed partial to complete occlusion of the alveolar air spaces by neutrophils and macrophages as well as thickening of the alveolar septa by similar inflammatory cells and edema, and some of the alveolar spaces exhibited protein (either fluid or fibrin). Histopathologic scores were significantly higher in Cl_2_ pigs compared to the air group (Fig. 5A). In Cl_2_ pigs, the tracheal mucosa was completely disrupted and overlaid with cell debris, whereas histopathologic analysis of the trachea from the air group showed normal architecture (Fig. 5B).

**Figure 5:**
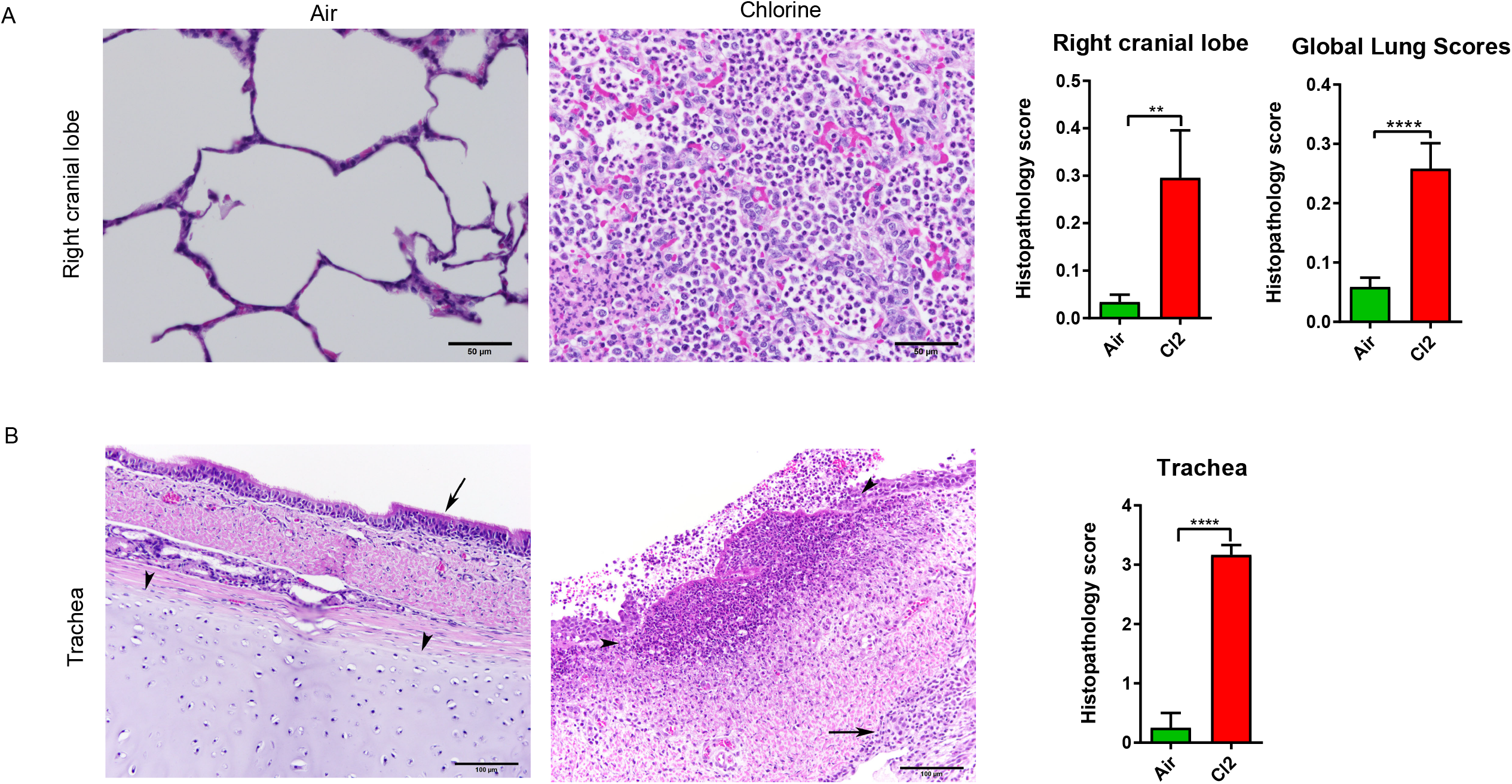
Histopathology of lungs in pigs exposed to chlorine gas. Anesthetized and mechanically ventilated pigs were exposed to either chlorine gas at ≤ 240 ppm or filtered room air for 1 hour. Histopathologic analysis was performed following *an official American Thoracic Society (ATS) Workshop Report:* Features and Measurements of Experimental Acute Lung Injury in Animals, with some modifications. **(Fig 5A)** Representative images of the right cranial lobe of lungs are presented. In the air group, normal open alveolar architecture of parenchyma is seen without any significant lesions. In the chlorine group, note partial to complete occlusion of the alveolar air spaces by neutrophils and macrophages, thickening of the alveolar septa by similar inflammatory cells, and edema. Some of the alveolar spaces exhibit protein (either fluid or fibrin). The bar graphs show histopathology scores both in the right cranial lobe and in all lobes. **(Fig 5B)** Representative tracheal images are presented. In the air group, normal trachea with pseudostratified columnar epithelium (arrows) and uniform lamina and cartilage (arrowheads) is presented. In the chlorine group, the tracheal mucosa is attenuated with the loss of cilia, and multifocal, abrupt ulcers and marked infiltrates by primarily neutrophils (suppuration) are present (outlined by arrowheads). The surface of the mucosa is overlaid by cell debris, and the deeper lamina (arrow) contains infiltrates by lymphocytes, plasma cells, and macrophages. Tracheal ring cartilage is visible in the air-exposed sections, but not in the chlorine-exposed sections due to edema and inflammation, although all images were taken at the same magnification. The bar graph shows the histopathology score of the trachea. **, *p* ≤ 0.01; ****, *p* < 0.0001. All images are presented at 40x magnification.

### Chlorinated fatty acids

Free and total (ie, free + esterified) 2-chlorofatty acids were detected in plasma and lung tissues from both groups. Free fatty acids (2-chloropalmitic acid, 16:0 Cl and 2-chlorostearic acid, 18:0 Cl), which are not esterified to complex lipids, were significantly increased in plasma samples collected within 2 hours after chlorine exposure (Figure 6A), whereas total fatty acids (16:0 Cl and 18:0 Cl) were significantly increased in plasma samples that were collected up to 24 hours after chlorine exposure compared to correspondingly timed samples from pigs in the air group (Figure 6B). Both free and total 2-chloropalmitic acid and 2-chlorostearic acid were significantly increased in lung tissue harvested at 24 hours after chlorine exposure compared to air exposure (Figure 6C-D).

**Figure 6:**
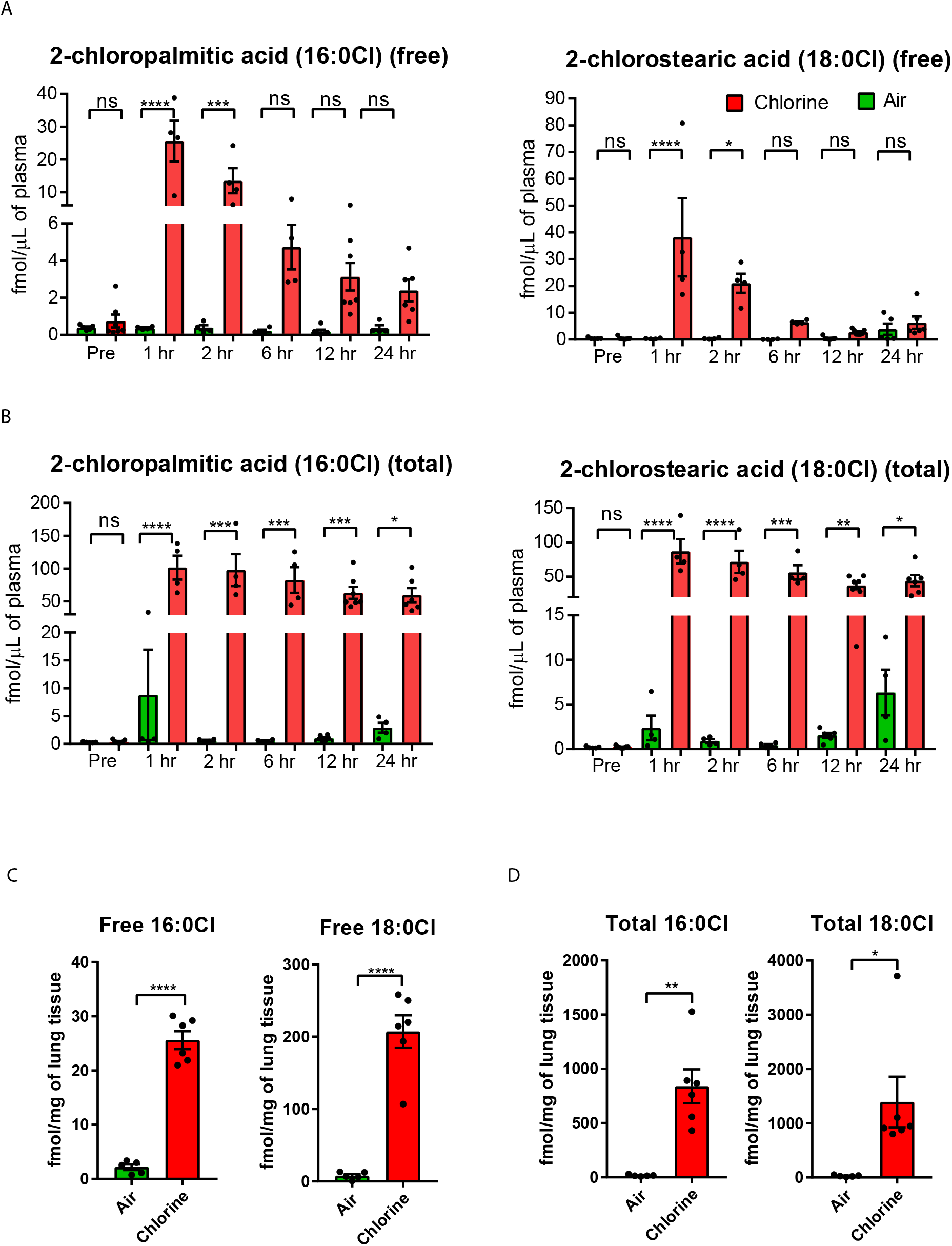
Chlorinated fatty acids in plasma and lung tissues of chlorine-exposed pigs. Free and total (ie, free + esterified) 2-chlorofatty acids were detected in plasma and lung tissues from both the chlorine and air groups. Free and total fatty acids (2-chloropalmitic acid, 16:0 Cl and 2-chlorostearic acid, 18:0 Cl) were increased in plasma samples that were collected at multiple time points until 24 hours post chlorine exposure (Figure 6A-B). Both free and total 2-chloropalmitic acid (Figure 6C) and 2-chlorostearic acid (Figure 6D) were significantly increased in lung tissue samples harvested at 24 hours post chlorine exposure vs filtered room air exposure. *, *p* ≤ 0.05; **, *p* ≤ 0.01; ***, *p* ≤ 0.001; ****, *p* < 0.0001; ns, not significant.

### Chlorinated tyrosine adducts

An analytical method to quantify Cl-Tyr and Cl_2_-Tyr in pig lung and plasma tissues was established (Table S2 and Figure S3). Concentrations of Cl-Tyr and Cl_2_-Tyr remained either close to lower reportable level (LRL) or undetectable in both plasma and lung tissue samples collected from air-exposed pigs whereas the concentrations have significantly increased in Cl_2_-exposed pigs by several folds (Figure 7).

**Figure 7:**
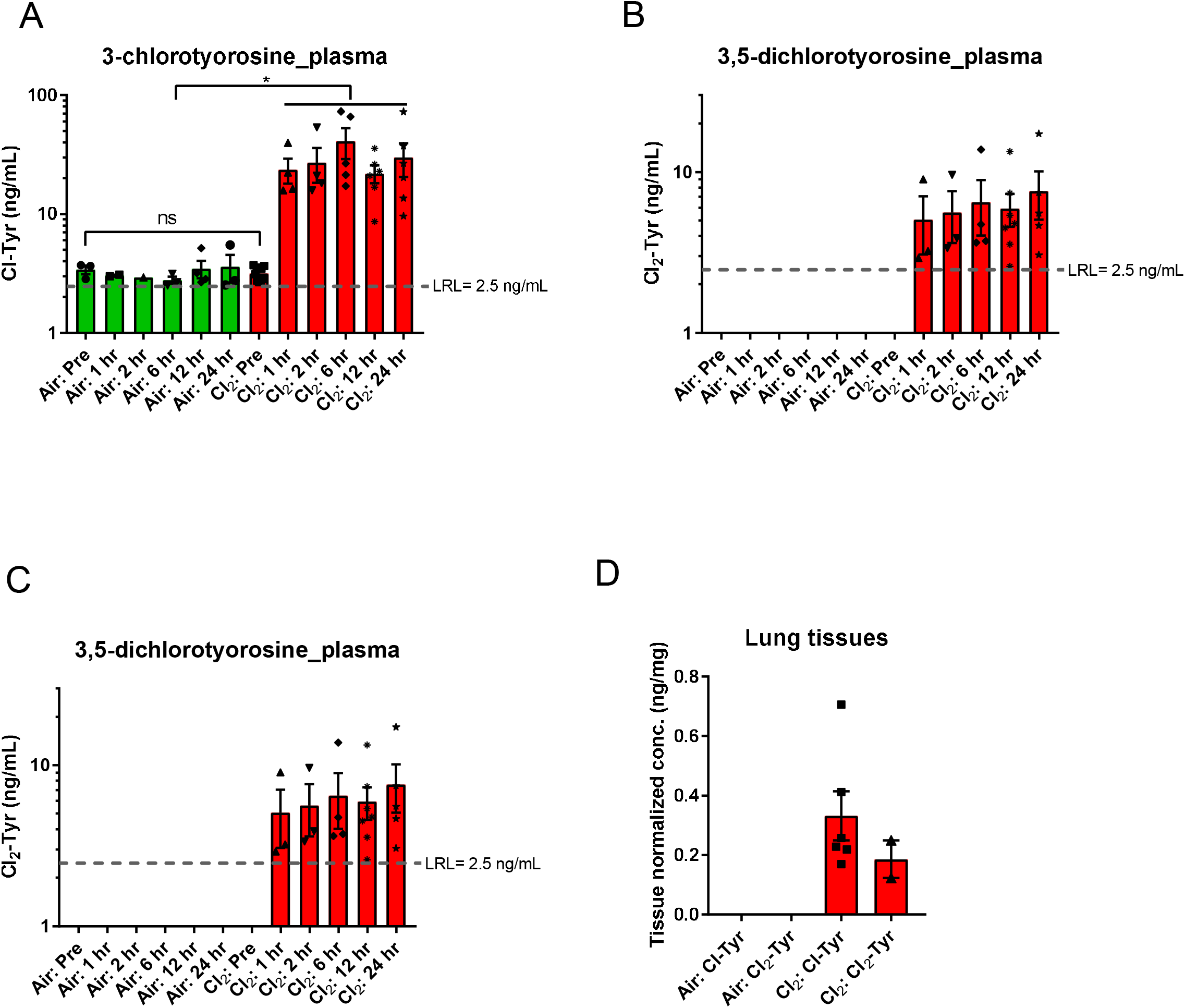
Chlorinated tyrosine adducts in plasma and lung tissues of chlorine-exposed pigs. 3-chlorotyrosine (Cl-Tyr) (Fig 7A) and 3,5-chlorotyrosine (Cl_2_-Tyr) (Fig 7B) concentrations remained close to lower reportable level (LRL) in plasma samples collected from air-exposed pigs whereas the concentrations have increased significantly in plasma samples collected from chlorine-exposed pigs. In air-exposed animals, the concentrations of Cl-Tyr and Cl_2_-Tyr were non-detectable whereas were increased by several folds in Cl_2_-exposed animals (Fig 7C-D). *p<0.05.

## DISCUSSION

In this study, we developed a human translational model of ALI in swine to meet the US FDA’s animal rule for the Chemical Medical Countermeasures Program [27]. The model described here overcomes several limitations described in the previously published literature [36-40]. The AUC of Cl_2_ exposure in this model was 12118 ± 584 ppm.min (mean ± SD), which correlates with fatal exposure in humans (> 400 ppm of Cl_2_ for 30 minutes, that is, 12000 ppm.min) [18]. Therefore, the swine model described in this study represents victims exposed to fatal concentrations of Cl_2_.

Although the literature review shows some reports on Cl_2_-induced ALI models in pig and sheep, those reports acknowledged limitations in their models [37-43]. We admit that no animal model is completely devoid of limitations. However, our model overcomes most of the limitations of previously published swine models. In contrast to the previously published models, we used a 5-cm H_2_O positive end-expiratory pressure (PEEP) to prevent the collapse of alveoli during exhalation, and to improve alveolar recruitment and oxygenation capacity [36-39]. While the accepted range of tidal volume (TV) is 6-8 mL/kg, previously published literature used TV outside the accepted volumes, ranging from 8-10 mL/kg [37], 10 mL/kg [39], 15-20 mL/kg [36], and 12 mL/kg [38], whereas we used 7 mL/kg. In previous studies, pigs were exposed to Cl_2_ at concentrations ranging from 100-140 ppm for 10 minutes (1000-1400 ppm.min) to 400 ppm for 15-20 minutes (6000-8000 ppm.min), whereas we used 240 ppm for 60 minutes (14400 ppm.min). We studied the pathophysiology of Cl_2_-induced ALI in pigs for 24 hours, whereas the available literature shows studies spanning over 5 hours. We maintained pigs on pressure-regulated volume control and assist control (PRVC A/C) mode of mechanical ventilation. In PRVC mode, the preset tidal volume is maintained by adapting to the changing compliance of the lungs to adjust inspiratory time and pressure. As assist control is also engaged in this setting, respiration is completely under the control of the mechanical ventilator.

### Acid-base imbalance

Previous clinical trials and pre-clinical studies suggested respiratory acidosis with exposure to Cl_2_, and further, intravenous or nebulized sodium bicarbonate (NaHCO_3_) has been used as a therapeutic agent to correct the acid-base imbalance. However, the evidence is mostly anecdotal, or at best, limited data are available to support this treatment [18, 44]. In the current study, we noted change acid-base imbalance from respiratory acidosis with metabolic acidosis initially after the end of Cl2 exposure and overall interpretation suggests respiratory alkalosis with metabolic acidosis. The increased lactate levels in Cl_2_-exposed pigs clearly suggest hypoxia and injury. The anesthesia protocol and mechanical ventilation may have had some influence on the acid-base imbalance. The lack of normal values for arterial blood gas analysis in pigs also makes it difficult to compare with human values. Therefore, the data should be interpreted with caution. Together, these findings suggest that generalized treatment of Cl_2_ victims with NaHCO_3_ may not be the Holy Grail, but still important for management of acid-base imbalance.

### Oxygenation parameters

Exposure to Cl_2_ resulted in immediate hypoxia and hypoxemia. The P/F ratio, also called the Carrico index, is a comparison between the oxygen level in the blood and the concentration of inhaled oxygen. The P/F ratio is generally used in critical care medicine to assess oxygenation in patients. PaO_2_/FiO_2_ ≤ 300 and ≤ 200 mmHg are signs of ALI and ARDS, respectively [45]. P/F ratio values had decreased below 300 in our chlorine group at the end of Cl_2_ exposure, and remained the same or decreased further over the following 24 hours. Oxygenation index (OI) is recognized as a more sensitive marker than the P/F ratio for assessing oxygenation because lung mechanics (mean airway pressure) is considered in the calculation.

Lower OI values indicate better utilization of inhaled oxygen, and are desirable. Our Cl_2_-exposed pigs had higher OI values. SpO_2_ indicates oxygen saturation levels in hemoglobin. We started all pigs at room air (FiO_2_ of 21%); however, we adjusted FiO_2_ to maintain at least 80% SpO_2_. The Cl_2_ group required higher FiO_2_ to maintain at least 80% SpO_2_ compared to the air group. As higher FiO_2_ values can result in the production of more reactive oxygen species, we tried to maintain a modest value of at least 80% SpO_2_. Targeting a higher SpO_2_ value by increasing FiO_2_ might be useful, but no controlled studies are available to support this idea, and supplementing higher FiO_2_ value has been linked to worsening ALI [46]. A-a gradient values were higher in the chlorine group vs the air group. Elevated A-a gradient values indicate that oxygen is not effectively transferred from the alveoli to the blood, and suggest a defect in diffusion and a ventilation-perfusion mismatch. This is not a surprising finding, as previous studies have shown that alveoli are damaged with Cl_2_ exposure.

### Respiratory physiologic parameters

Airway resistance and PIP increased significantly in Cl_2_-exposed pigs. C_dyn_ decreased rapidly following Cl_2_ exposure. This rapid decrease in C_dyn_ may be due to increased resistance to airflow, resulting in regional trapping of air with hyperinflation of lung regions distal to the site of bronchoconstriction. The collapse of pulmonary parenchyma and increase in interstitial fluids may also decrease C_dyn_ at later stages of ALI [38, 47]. In the methacholine airway challenge test, control animals tolerated higher concentrations of methacholine, suggesting airway hyperreactivity in Cl_2_-exposed animals. Further, there was a clear difference in baseline values between the 2 groups before administration of the first dose of methacholine, which is commonly not noted in rodent models of ALI.

### Pro-inflammatory cytokine markers

Pro-inflammatory cytokine markers, such as IL-6 and VEGF, were increased in the chlorine group compared to control animals. IL-6 values were consistently increased after Cl_2_ exposure in all 3 biological matrices tested. Therefore, IL-6 may be useful as a potential pro-inflammatory cytokine marker to assess disease progression as well as therapeutic efficacy of drug candidates.

### Effects on cardiovascular function

One pig died during Cl_2_ exposure, and another pig died 3-4 hours after Cl_2_ exposure. Both died rapidly without displaying any symptoms except progressive non-responsive hypotension, decreased heart rate, and hypoxemia. Cardiopulmonary resuscitation (CPR) efforts with 100% oxygen supplementation, epinephrine injection and defibrillation were not successful. A postmortem examination revealed that both pigs had pre-existing left ventricular hypertrophy (right-to-left ventricular wall thickness ratio ≥ 1:5). Previous studies show that cardiac ventricular hypertrophy is a common finding in pigs that is based on the birth season, breed, sex, and sire family [48, 49]. This suggests that Cl_2_ exposure in the context of pre-existing cardiac dysfunction can result in severe casualties. Cl_2_ exposure decreases cardiac output probably due to right ventricular failure secondary to pulmonary vasoconstriction [38]. Often, severe right ventricular failure results in death, particularly in patients with pre-existing left ventricular failure or left ventricular hypertrophy. Limited animal studies and isolated reports in humans exposed to Cl_2_ suggest effects on the cardiovascular system [50]. Additional detailed studies on cardiovascular effects are warranted.

### Postmortem lung gross examination and histopathology

Gross examination of the lungs revealed diffuse atelectatic lesions in the diaphragmatic lobes of both groups. These findings are consistent with the lesions commonly seen in animals restrained in dorsal recumbency (supine positioning) and mechanically ventilated. In this study, animals were mechanically ventilated in the supine position, which is one of the limitations of the study. Prone positioning was shown to be advantageous over supine positioning in both human studies and translational animal models [38, 51-53]. Future studies are warranted to prospectively study the advantages of prone positioning and positional maneuvers in Cl_2_-exposed pigs. The Cl_2_-exposed lung tissues exhibited all 3 key features of human ALI: neutrophilic alveolitis, deposition of hyaline membranes, and formation of microthrombi. Therefore, this animal model can be used to study other forms of ALI in humans [30, 31].

### CFAs and CTAs

There are no diagnostic markers for Cl_2_-exposed victims. The current literature on diagnostic markers is limited to pre-clinical studies, particularly in rodent models or in vitro studies [3-6, 34]. Free and total CFAs (2-chloropalmititc acid and 2-chlorostearic acid) were significantly increased in both plasma and lung tissues collected from the Cl_2_ group, suggesting that they may be used as diagnostic markers. Both free and total 2-chloropalmitic and 2-chlorostearic acids were significantly increased immediately after exposure to Cl_2_, and decreased over the following 24 hours. However, the levels remained several-fold higher until 24 hours after Cl_2_ exposure. We expect that CFA levels remain higher by several fold until 72 hours post exposure or beyond, as seen in rodent studies [3]. These chlorinated lipids are produced when Cl_2_ or hypochlorite reacts with plasmalogen. Previously, in rodent models of Cl_2_-induced ALI, chlorinated lipids such as 2-chloropalmitaldehyde (2-Cl-Pald), 2-chlorostearaldehyde (2-Cl-Sald), and their oxidized products, free- and esterified 2-chloropalmitic acid (2-Cl-PA) and 2-chlorostearic acid were detected in the lungs and plasma [3, 54]. Previously CTAs (Cl-Tyr and Cl2-Tyr) have been detected in various mouse biological matrices, and in human blood samples under in vitro conditions [34, 35]. Overall, Cl-Tyr and Cl_2_-Tyr concentrations were undetectable or remained below lower reportable level (LRL) in both plasma and lung tissue samples collected from air-exposed pigs whereas the concentrations were detectable or several folds higher in plasma and lung tissue samples collected from chlorine-exposed pigs. Thus, the findings in plasma and lung tissues of our porcine model corroborate with previous findings in rodents. Detection of CFAs and CTAs independently in 2 animal models of Cl_2_-induced ALI strongly suggests their utility as potential biomarkers. Particularly, detection of CFAs and CTAs in plasma samples until 24 hours is advantageous from a forensic point of view, and the ease of sample collection at multiple time points is also advantageous. Detection of CFAs and CTAs in lung tissue samples is also highly relevant from a forensic point of view in establishing that victims indeed died of Cl_2_ exposure.

### Clinical applicability

The actual pathophysiologic course of human victims exposed to Cl_2_ is highly variable depending on the concentration of the gas, duration of exposure, and distance of the individual victim from the locus of the source. This model represents several features of humans exposed to fatal concentrations of Cl_2_. When a human is exposed to Cl_2_, the victim runs, and under those horrified panic situations, the victim breathes through the oropharynx. Several diffusion modeling studies at low Cl_2_ concentrations suggest that ∼ 90% of inhaled Cl_2_ will be scrubbed in the nasal passages and pharynx, and that only 10% of inhaled Cl_2_ reaches beyond the hypopharynx [55]. In our model, pigs were exposed to Cl_2_ via an endotracheal tube not only to ensure homogenous exposure but also to replicate human Cl_2_ exposure.

### Limitations of the study

We limited our study observation time to 24 hours. However, in the continuum of this work, it would be worthwhile to extubate the animals after Cl_2_ exposure, and monitor them over extended time, at least up to 2 months. Animals under continued mechanical ventilation and anesthesia fail to exhibit their natural physiologic protective reflexes such as cough, apnea, and shallow breathing [38]. Therefore, studies are warranted to understand the long-term pathophysiology of Cl_2_-exposed and extubated pigs.

## CONCLUSIONS

We recapitulated the pathophysiologic course of Cl_2_ on the pulmonary system in pigs, and demonstrated that our pig model of acute lung injury serves as an excellent translational model of human Cl_2_-induced ALI for developing/screening potential chemical countermeasures. Further, we identified biomarkers of chlorine exposure.

## Acknowledgments

Authors would like to thank Dr Michael Foster, PhD, for his thoughtful discussions in setting up our chlorine exposure system; Ianthia E Parker, George Quick, Michael Lowe, and Christie Holmes for their assistance in pig studies; Dr Jerry Ritchey, DVM, PhD, DACVP, Center for Veterinary Health Sciences, Oklahoma State University, for blinded histopathologic analysis.

## AUTHOR CONTRIBUTIONS

SA and S-EJ conceived and designed research; SA and MG performed animal experiments; CJA, KAS, DAF, and RPP performed chlorinated fatty acid analyses; BSC, JQG, JWP, and TAB performed chlorinated tyrosine adduct analyses; SA and S-EJ analyzed data and interpreted experimental results; SA prepared figures and wrote the first draft of manuscript; SA and S-EJ edited and revised manuscript; SA and S-EJ approved final version of manuscript.

