## supplementary information for "Recapitulation of Human Pathophysiology and Identification of Forensic Biomarkers in a Translational Swine Model of Chlorine Inhalation Injury"

**MATERIALS AND METHODS**

***Preparation of animals and anesthesia:*** Specific pathogen-free (SPF) male and female Yorkshire pigs (n = 14) (castrated males and intact females) (30-40 kg) were procured, and given at least 48 hours to acclimate. Pigs were housed in an Association for Assessment and Accreditation of Laboratory Animal Care (AAALAC) International-accredited facility at Duke University School of Medicine, Durham, NC, USA. All animal husbandry and study procedures in this study were approved by the Duke University Institutional Animal Care and Use Committee (IACUC). Pigs were fasted overnight before the study with *ad libitum* access to water. Animals were pre-medicated with acepromazine (1.1 mg/kg) and ketamine (22 mg/kg) intramuscularly. Anesthesia was induced with a bolus dose of propofol (5-10 mg/kg iv) titrated to the effect, and then maintained with continuous rate infusion (CRI) of propofol at 5-10 mg/kg/h. Fentanyl was given as CRI at the rate of 20-40 µg/kg/h for analgesia. Animals were intubated with a 7.5-mm ID endotracheal tube, and connected to a mechanical ventilator (AVEA™ ventilator, Viasys Healthcare Critical, Palm Springs, CA, USA) with the following settings: pressure-regulated volume control and assist control (PRVC A/C) mode; tidal volume: 7 mL/kg; respiratory rate: 25 bpm; inspiratory time: 0.7 seconds; Positive End Expiratory Pressure (PEEP): 5 cm H_2_O; inspiratory fraction of oxygen (FiO_2_): 21%, adjusted to maintain SpO_2_ levels above 80% to provide adequate tissue oxygenation. An airway adapter was placed in line between the endotracheal tube and the ventilator circuit to monitor respiratory physiologic parameters. Tidal volume, respiratory rate (RR), dead space, peak airway pressure, VCO_2_, mean airway pressure (mPaw), airway resistance (Raw), end-tidal carbon dioxide, and dynamic lung compliance (C_dyn_) were measured on a breath-to-breath basis using respiratory profile monitor NM3 (Respironics NM3, Philips Healthcare, Wallingford, CT, USA). Femoral artery and vein were surgically catheterized for arterial pressure monitoring and intravenous fluid administration, respectively. A Foley catheter was placed into the bladder with a standard cystotomy surgical procedure. Heart rate, EKG, end-tidal CO_2_, SpO_2_, respiratory rate, and body temperature were continuously monitored. Pigs were given a CRI of normal saline at 10 mL/kg/h (bolus dose) initially and then at 5 mL/kg/h (maintenance dose). Arterial blood gas (ABG) analyses were performed at hourly intervals using Gem Premier 3000, Model 5700 (Instrumentation Laboratory, Lexington, MA, USA). Isothermia was maintained with water-heating/cooling blankets and insulation blankets. Complete blood counts (CBC) and serum chemistry panels were assessed using IDEXX ProCyte Dx Hematology Analyzer (IDEXX Laboratories, Inc, Westbrook, Maine, USA) and Heska Dry Chem 7000 (Heska, Inc, Loveland, CO, USA), respectively, at baseline and at 24 hours after air or chlorine gas exposure.

***Chlorine gas exposure:*** After surgical instrumentation and acquisition of baseline data (respiratory and cardiovascular physiologic parameters, and arterial blood gas analysis), animals were randomized to chlorine gas exposure (n = 8) or filtered room air exposure (n = 6). However, 2 pigs died during or a few hours after exposure to chlorine gas. Therefore, data were presented for 6 pigs in the chlorine group. Chlorine gas was delivered via the inspiratory limb of the mechanical ventilator circuit, and was monitored with a chlorine detector (Porta Sense II Gas Leak Detector, AFC International, Inc., DeMotte, IN, USA) at 8-15-minute intervals to ensure delivery of ≤ 240 ppm for 1 hour. Pigs were exposed to chlorine gas in a negatively pressurized room with adequate ventilation and monitors for leak detection in multiple locations. Chlorine gas exhaust from the mechanical ventilator, chlorine detectors, and any potential leak in the custom-made chlorine entrapment chamber was scavenged through a series of canisters with impingers containing a basic solution, water, and acid indicator, and then vented through a dedicated exhaust system. Figures 1A and 1B show the study paradigm and the schematic of the chlorine gas exposure system. The key respiratory physiologic parameters were recorded and arterial blood gas analysis was performed at 5 and 10 minutes post chlorine gas exposure, and then subsequently at 1-hour intervals until euthanasia at 24 hours post chlorine gas exposure.

***Oxygenation parameters:*** The ratio of the partial pressure of oxygen in arterial blood and fraction of inhaled oxygen (PaO_2_/FiO_2_), oxygenation index, SpO_2_/FiO_2_, and arterial-alveolar (A-a) gradient were the oxygenation parameters primarily considered. The ratio of the partial pressure of oxygen in arterial blood to the fraction of inhaled oxygen is a comparison between the oxygen level in arterial blood and the O_2_ concentration that is breathed. PaO_2_/FiO_2_ ≤ 300 and ≤ 200 mmHg are signs of ALI and ARDS, respectively. Oxygenation index (OI) is calculated using the following formula:

Oxygenation index = (FiO_2_ x mPaw) x100 /PaO_2_

*Where FiO_2_ = the fraction of oxygen in the inhaled gas*

*mPaw = mean airway pressure*

*PaO_2_ = the partial pressure of oxygen in arterial blood*

Arterial-alveolar (A-a) gradient was determined using the following formula:

A-a gradient = PAO_2_ – PaO_2_

PAO_2_ = (P_atm_ – P_water_) FiO_2_ – (PaCO_2_/0.8)

*Where PAO_2_ = the partial pressure of oxygen in alveoli*

*P_atm_ = the atmospheric pressure (760 mm Hg, at sea level)*

*P_water_ = the vapor pressure of water at body temperature (47 mm Hg, at sea level)*

*FiO_2_ = the fraction of O_2_ in the inhaled gas*

*PaCO_2_ = the partial pressure of CO_2_ in arterial blood*

*0.8 = value used for the respiratory quotient*

*PaO_2_ = the partial pressure of O_2_ in arterial blood*

***Respiratory physiologic parameters:*** Compliance dynamics (C_dyn_), airway resistance (Raw), and peak inspiratory pressure (PIP) were analyzed at baseline and at 5 and 10 minutes post chlorine gas exposure, and then hourly for 24 hours.

***Methacholine airway challenge test:*** The airway mechanics were measured by challenging pigs to increasing concentrations of methacholine hydrochloride (0, 0.5, 1, 2, 4, 8, 16, and 32 mg/mL) (Sigma-Aldrich, St. Louis, MO, USA) via an Aeroneb™ nebulizer (Aerogen Ltd, Ireland) at 24 hours post chlorine gas or air exposure.

***Bronchoscopy and bronchoalveolar lavage fluid collection:*** Bronchoscopy was performed just prior to euthanasia. A 22 French bronchoscope was inserted into the endotracheal tube while the animal was maintained under anesthesia. The scope was placed into the right primary bronchus. Thirty mL of phosphate buffered saline (PBS) solution containing 0.1% BSA and protease inhibitors (Roche, Indianapolis, IN, USA) was flushed through the bronchoscope and retrieved immediately into a sterile vial. This procedure was repeated one more time, and the bronchoalveolar lavage fluid (BALF) was saved on ice for differential leukocyte counting using cytospin procedures.

***Postmortem examination and histopathology analysis:*** At the end of the 24-hour period post gas exposure, pigs were euthanized using an IACUC-approved protocol that followed *AVMA guidelines on euthanasia*. Lungs were removed from pigs for gross pathologic examination. For pro-inflammatory cytokine and diagnostic marker analyses, lung tissues were flash frozen in liquid nitrogen, and then stored at -80^0^C. For histopathologic analysis, lung specimens were collected in 10% formalin in neutral buffered saline. Lung biopsy samples were processed for histopathology examination at Oklahoma Animal Disease Diagnostic Laboratory, Center for Veterinary Health Sciences, Stillwater, OK. Briefly, lung tissues were embedded in paraffin, sectioned at 5-µm thickness, and stained with hematoxylin and eosin (H&E). A board-certified veterinary pathologist performed blinded histopathologic scoring of lung specimens following *an official American Thoracic Society (ATS) Workshop Report:* Features and Measurements of Experimental Acute Lung Injury in Animals, with some modifications [1]. Briefly, 4 specimens (cranial and caudal lobes of both right and left lungs) from each animal were observed, and 10 high-power fields (40X) in each lung specimen were assessed for neutrophils in the alveolar space, neutrophils in the interstitial space, protein in alveoli, hyaline membranes, and alveolar septal thickening, ie, 40 high magnification fields per animal. Scores were given per each observed high magnification field (Table S1, adapted from [1]).

**Table S1. The scoring method used for the histopathologic analysis of lung biopsy samples.** Histopathologic scoring of lung specimens was performed following *an official American Thoracic Society (ATS) Workshop Report:* Features and Measurements of Experimental Acute Lung Injury in Animals, with some modifications. The table was adapted from Matute-Bello et al, 2011. Briefly, 10 high-power fields (40X) for each of 4 lung specimens/animal were assessed for neutrophils in the alveolar space, neutrophils in the interstitial space, protein in alveoli, hyaline membranes, and alveolar septal thickening. A score for each of these parameters was assigned to every high magnification field.

| **Parameter** | | **Score per field** | | |
| --- | --- | --- | --- | --- |
|  |  | 0 | 1 | 2 |
| A | Neutrophils in the alveolar space | None | 1-5 | >5 |
| B | Neutrophils in the interstitial space | None | 1-5 | >5 |
| C | Hyaline membranes | None | 1 | >1 |
| D | Proteinaceous debris filling the air spaces | None | 1 | >1 |
| E | Alveolar septal thickening | <2x | 2x-4x | >4x |

The scores for each parameter were then used to calculate the ALI score for each specimen using the following equation:

Score = [(20 x A) + (14 x B) + (7 x C) + (7 x D) + (2 x E)] / (number of fields x 100)

(We observed 10 high magnification fields per slide/tissue specimen instead of 20 high magnification fields as suggested in the ATS guidelines. However, tissue specimens were collected from different regions to accommodate the heterogeneous injury pattern noted in this model. Altogether, 10 high magnification fields from each of 4 different tissues/animal, ie, 40 high magnification fields/animal, were evaluated in this study.)

Chlorine gas has intermediate water solubility, and results in injury of upper and lower airways. The trachea is the first site of exposure to chlorine gas in the current model. Since we also observed gross pathologic lesions in the trachea with chlorine exposure, we developed a new method for scoring histopathologic lesions in the trachea on a severity scale of 0-4 (0 = no microscopic lesions; 1 = mild lesions; 2 = moderate lesions; 3 = severe lesions; 4 = chronicity with re-epithelialization and early fibroplasia).

***Lung wet/dry weight ratio***: The lung wet/dry weight ratio was used as a surrogate marker for edema. To determine the wet/dry lung weight ratio, 4 small specimens from the cranial and caudal lobes of the right and left lungs were weighed (wet weight), and then incubated at 50^0^C for 48 hours and weighed again (dry weight).

***Total and differential BALF leucocyte counts:*** Total and differential leukocyte counts were obtained following previously published methods, with some modifications [2]. Briefly, BALF samples were centrifuged, and the supernatant was saved for analyses of pro-inflammatory cytokine markers and diagnostic markers. The pellet was re-suspended in 10 mL of RBC lysis buffer (1x) (BD Biosciences, East Rutherford, NJ, USA), and incubated at room temperature for 10-15 minutes. RBC lysis was terminated by adding an equal volume of HBSS (1x) (Gibco, Life Technologies, Grand Island, NY), and the suspension was centrifuged again. The supernatant was discarded, and the pellet was re-suspended in 1 mL of HBSS. The suspension was further diluted 1:50 or higher, and the diluted cells were counted for the total leukocyte counts using a hand-held cell counter (Scepter^®^, Millipore, Billerica, MA, USA). Cytospin and Diff-Quick staining procedures were performed following standard procedures [2].

***Quantification of pro-inflammatory cytokine markers in BALF, serum, and lung tissues:*** The concentrations of pro-inflammatory cytokine markers were determined in BALF supernatants, serum samples, and lung tissue homogenates using ELISA kits (R&D Systems, Minneapolis, MN, USA) per manufacturer’s instructions. Each sample was assayed in duplicate. To homogenize lung tissue, 1 mL of phosphate buffered saline with protease inhibitors (Roche) was added, and the sample was homogenized using a hard tissue disrupter attached to a handheld homogenizer (Omni International, Kennesaw, GA, USA) [3]. The data were analyzed using a standard curve.

***Chlorinated fatty acids in lungs and plasma:*** Free and total (ie, free + esterified) 2-chlorofatty acids were measured as previously described by LC/MS, following Dole extraction [4]. Total 2-chlorofatty acids were measured by LC/MS after base hydrolysis, while free 2-chlorofatty acids were not subjected to base hydrolysis. Extractions were performed using 25 µL of plasma spiked with 103.5 fmol of 2-chloro-[d4-7,7,8,8]-palmitic acid (2-[d4]-ClPA) as the internal standard. Lung tissue (20 mg) was pulverized and subjected to Bligh-Dyer lipid extraction [5] in the presence of 103.5 fmol of 2-[d4]-ClPA (internal standard). Half of the lung lipid extract was then subjected to LC/MS analyses for free 2-chlorofatty acids, and the other half of the extract subjected to base hydrolyses followed by LC/MS analyses for total 2-chlorofatty acids.

***Chlorinated tyrosine adducts in lungs and plasma***: The biomarkers 3-chlorotyrosine (Cl-Tyr) and 3,5-dichlorotyrosine (Cl2-Tyr) were in plasma and lung tissues samples from air or chlorine exposed pigs. Materials were purchased from Fisher Scientific, Sigma-Aldrich, MG Scientific, Eppendorf, Cambridge Isotopic Laboratories, IsoSciences, Omni International and Tennessee Blood Services as described by Pantazides *et al.* [6].

*Preparation of calibrators and QCs***:** 3-chloro-L-tyrosine and 3,5-dichloro-L-tyrosine were used to prepare seven calibrators ranging from 2.5-500 ng/mL in 0.1% formic acid in water. Internal standard containing ^13^C_6_-isotopically labeled 3-chloro-L-tyrosine and ^13^C_9,_^15^N-isotopically labeled 3,5-dichloro-L-tyrosine was prepared at 25.0 ng/mL in 0.1% formic acid in water. Three quality control materials were made by adding sodium hypochlorite to plasma to create a QC low, QC mid, and QC high.

*Sample preparation:* Lung tissue preparation was previously described by Pantazides *et al. [6]*. In brief, a portion of the lung tissue (~5 mg) was transferred to a 2 mL vial containing 50 mM ammonium bicarbonate (200 uL) and metal grinding beads. The vial was loaded onto a Omni Bead Ruptor Elite tissue homogenizer for 3 cycles of 50 sec at a speed of 5.5 m/s. Using these settings, the lung tissue was uniformly homogenized. Chlorinated tyrosine was extracted from plasma and homogenized lung tissue samples as described by Crow and Pantazides *et al.* [6, 7]. In brief, plasma and homogenized lung tissue, calibrators, and QCs (10 µL) were added to a 2 mL 96-well plate. ISTD (20 µL) and 2 mg/mL pronase in 50 mM ammonium bicarbonate (100 µL) were added to each well. The plate was sealed and shaken at 700 rpm for 75 min at 60^o^C. After incubation, each sample was added to a 96-well protein precipitation plate containing acetonitrile (360 µL) and eluted into a new 96-well plate using a vacuum manifold. Samples were dried using a nitrogen stream, resuspended in 0.1% formic acid in water (50 µL), transferred to a 96-well PCR plate, and analyzed via LC-MS/MS.

*UHPLC-MS/MS – instrument settings and optimization:* Instrument settings were previously described by Pantazides *et al.* [6]. Cl-Tyr and Cl_2_-Tyr were detected on an Agilent 6490 triple quadruple mass spectrometer coupled to the Agilent 1290 Infinity II UHPLC system. The UHPLC system contained a sampler, binary pump, and column compartment. Analytical separation was performed on a Hypercarb column (2.1 x 30mm, 3 µm) heated to 60^o^C. Mobile phase A (MPA) was 0.1% formic acid in water and mobile phase B (MPB) was 0.1% formic acid in acetonitrile. The gradient started with a 2% MPB hold from 0 to 1 min; gradual increase in MPB from 2% to 50% from 1.01 to 4 min; a step to 98% MPB from 4.01 to 4.50 min; and ended with a step down to 2% MPB at 4.51 min using a consistent flow rate of 250 µL/min. **Table S2** shows ESI-MS/MS parameters for detection of native and labeled chlorotyrosines**.** The ESI-MS/MS parameters were described by Pantazides *et al.* [6]. The source operated in positive mode with a gas temp of 290^o^C, gas flow of 11 L/min, nebulizer of 60 psi, sheath gas temperature of 200°C, sheath gas flow of 11.0 L/min, drying gas of 14.00 L/min, capillary voltage of 1000 V, and nozzle voltage of 300 V.

|  | **QQQ Acquistion** | | | | |
| --- | --- | --- | --- | --- | --- |
| **Parameter** | **Transition** | **Dwell Time (ms)** | **Fragmentor (V)** | **Collision Energy (V)** | **Cell Accelerator Voltage (V)** |
| 3,5-dichlorotyrosine (Quant) | 250.0 → 169.0 | 20 | 380 | 30 | 2 |
| 3,5-dichlorotyrosine (Conf) | 250.0 → 204.0 | 20 | 380 | 10 | 2 |
| 3,5-dichlorotyrosine (ISTD) | 260.0 → 178.0 | 20 | 380 | 30 | 2 |
| 3-chlorotyrosine (Quant) | 216.0 → 135.1 | 20 | 380 | 30 | 2 |
| 3-chlorotyrosine (Conf) | 216.0 → 170.0 | 20 | 380 | 10 | 2 |
| 3-chlorotyrosine (ISTD) | 222.0 → 205.0 | 20 | 380 | 5 | 2 |

*Method validation:* Matrix effects, extraction recovery, sensitivity, specificity, calibration curve, linearity, limit of detection, limit of quantitation, lowest reportable limit,and intra- and inter-day precision and accuracies were described in detail by Crow *et al.* and Pantazides *et al. [6, 7]* for plasma and lung tissue samples, respectively. Figure S3 shows overlay of extracted ion chromatograms for plasma and lung samples collected from pigs pre- and post-chlorine gas exposure.

**Table S3.** Mean complete blood counts (CBC) of pigs exposed to filtered air or chlorine gas

| **Parameter** | **Units** | **Baseline** | | | | **24 hours post exposure** | | | |
| --- | --- | --- | --- | --- | --- | --- | --- | --- | --- |
|  |  | **Air** | | **Cl2** | | **Air** | | **Cl2** | |
|  |  | Mean | SD | Mean | SD | Mean | SD | Mean | SD |
| RBC | M/µL | 5.2 | 0.36 | 5.5 | 0.46 | 4.2 | 0.45 | 5.0 | 1.03 |
| HGB | g/dL | 9.3 | 0.87 | 9.1 | 0.62 | 11.6 | 1.74 | 10.3 | 2.49 |
| HCT | % | 28.0 | 2.12 | 27.2 | 1.68 | 24.4 | 2.99 | 24.6 | 4.39 |
| MCV | fL | 54.8 | 0.87 | 49.3 | 2.20 | 56.6 | 2.87 | 49.2 | 2.47 |
| MCH | pg | 17.9 | 0.63 | 16.4 | 0.55 | 26.9 | 2.03 | 20.6 | 3.69 |
| MCHC | g/dL | 33.0 | 0.86 | 33.3 | 0.77 | 47.6 | 2.86 | 42.1 | 8.54 |
| RDW-SD | fL | 35.3 | 2.61 | 34.6 | 1.34 | 37.2 | 4.81 | 33.8 | 1.44 |
| RDW-CV | % | 20.3 | 1.61 | 22.6 | 2.15 | 19.8 | 1.82 | 21.8 | 1.82 |
| RET | K/µL | 56.3 | 26.46 | 65.9 | 39.80 | 42.6 | 19.75 | 74.4 | 44.58 |
| RET | % | 1.1 | 0.55 | 1.2 | 0.78 | 1.0 | 0.50 | 1.6 | 1.05 |
| PLT | K/µL | 335.2 | 115.59 | 265.8 | 24.23 | 208.0*^a^* | 90.02 | 127.6*^b,c^* | 61.44 |
| PDW | fL | 9.8 | 1.65 | 11.3 | 3.89 | 9.6 | 0.00 |  |  |
| MPV | fL | 7.5 | 1.04 | 7.8 | 1.56 | 7.4 | 0.00 |  |  |
| P-LCR | % | 11.9 | 6.36 | 13.0 | 11.24 | 11.8 | 0.00 |  |  |
| PCT | % | 0.3 | 0.06 | 0.2 | 0.06 | 0.2 | 0.00 |  |  |
| WBC | K/µL | 16.3 | 4.54 | 16.5 | 1.50 | 14.2 | 5.32 | 12.3 | 2.03 |
| NEUT | K/µL | 8.1 | 3.65 | 5.4 | 2.19 | 6.7 | 3.35 | 3.6 | 2.38 |
| LYMPH | K/µL | 7.4 | 0.93 | 10.0 | 1.63 | 4.7 | 1.75 | 6.5 | 2.36 |
| MONO | K/µL | 0.8 | 0.53 | 1.0 | 0.48 | 0.7 | 0.28 | 0.6 | 0.19 |
| EO | K/µL | 0.0 | 0.02 | 0.1 | 0.07 | 1.3 | 0.98 | 1.4 | 2.13 |
| BASO | K/µL | 0.03 | 0.01 | 0.03 | 0.02 | 0.8 | 1.19 | 0.3 | 0.57 |
| NEUT | % | 47.1 | 12.58 | 32.3 | 12.25 | 45.5 | 8.95 | 28.1 | 16.46 |
| LYMPH | % | 47.6 | 11.31 | 61.2 | 12.52 | 34.2 | 7.24 | 55.3 | 24.02 |
| MONO | % | 4.8 | 2.52 | 5.8 | 2.42 | 4.7 | 1.84 | 4.7 | 1.90 |
| EO | % | 0.3 | 0.19 | 0.6 | 0.47 | 9.7 | 7.95 | 10.0 | 13.75 |
| BASO | % | 0.2 | 0.05 | 0.2 | 0.11 | 5.9 | 7.27 | 1.8 | 3.69 |

Data are presented as mean ± SD (n = 5 per group). A significant difference between groups was determined by the Holm-Sidak method (*t*-test), with alpha = 0.05. Computations assume that all samples are from populations with the same scatter (SD). Significant differences between values are indicated by: *^a^,* baseline and 24-hour time points in the air group; *^b^*, baseline and 24-hour time points in the chlorine group; and *^c^*, 24-hour time points in the air and chlorine groups.

RBC = red blood cells; HGB = hemoglobin; HCT = hematocrit; MCV = mean corpuscular volume; MCH = mean corpuscular hemoglobin; MCHC = mean corpuscular hemoglobin concentration; RDW-SD = red blood cell distribution width standard deviation; RDW-CV = red blood cell distribution width coefficient of variation; RET = reticulocytes; PLT = platelets; PDW = platelet distribution width; MPV = mean platelet volume; P-LCR = platelet larger cell ratio; PCT = plateletcrit; WBC = white blood cells; NEUT = neutrophils; LYMPH = lymphocytes; MONO = monocytes; EO = eosinophils; BASO = basophils.

**Table S4.** Serum chemistry panel of pigs exposed to filtered air or chlorine gas

| **Parameter** | **Unit** | **Baseline** | | | | **24 hours post exposure** | | | |
| --- | --- | --- | --- | --- | --- | --- | --- | --- | --- |
|  |  | **Air** | | **Cl2** | | **Air** | | **Cl2** | |
|  |  | Mean | SD | Mean | SD | Mean | SD | Mean | SD |
| BUN | mg/dL | 8.9 | 2.08 | 9 | 2.38 | 11.0 | 4.03 | 12.3 | 5.14 |
| Creatinine | mg/dL | 1.0 | 0.12 | 1 | 0.13 | 0.9 | 0.18 | 1.1 | 0.32 |
| BUN/Creatinine Ratio |  | 8.9 | 1.09 | 9 | 3.09 | 11.6 | 2.84 | 10.9 | 1.59 |
| Phosphorus | mg/dL | 10.9 | 0.77 | 9 | 1.83 | 10.9 | 0.68 | 10.3 | 1.50 |
| Calcium | mg/dL | 8.7 | 0.55 | 9 | 0.48 | 7.5 | 0.51 | 7.5 | 0.69 |
| Total protein | g/dL | 5.0 | 0.40 | 5 | 0.51 | 4.4 | 0.16 | 4.3 | 0.56 |
| Albumin | g/dL | 2.9 | 0.33 | 3 | 0.17 | 2.0 | 0.26 | 1.9 | 0.18 |
| Globulin | g/dL | 2.1 | 0.24 | 2 | 0.39 | 2.4 | 0.31 | 2.4 | 0.42 |
| Alb/Glob ratio |  | 1.4 | 0.23 | 1 | 0.15 | 0.8 | 0.18 | 0.8 | 0.13 |
| Glucose | mg/dL | 110.0 | 18.72 | 120 | 48.97 | 61.8*^a^* | 16.48 | 56.4*^b^* | 22.88 |
| Cholesterol | mg/dL | 78.2 | 16.66 | 76 | 24.53 | 151.0*^a^* | 29.97 | 116.6*^b^* | 28.88 |
| ALT (GPT) | U/L | 41.6 | 10.81 | 44 | 16.24 | 44.6 | 13.58 | 53.4 | 15.21 |
| ALP | U/L | 167.6 | 54.98 | 152 | 41.31 | 231.2*^a^* | 91.50 | 210.6*^b^* | 75.30 |
| GGT | U/L | 22.6 | 3.71 | 37 | 8.94 | 26.0 | 2.55 | 31.4 | 4.04 |
| Total Bilirubin | mg/dL | 0.1 | 0.04 | 0 | 0.05 | 0.2 | 0.09 | 0.2 | 0.05 |

Data are presented as mean ± SD (n = 5 per group). A significant difference between groups was determined by the Holm-Sidak method (*t*-test), with alpha = 0.05. Computations assume that all samples are from populations with the same scatter (SD). Significant differences between values are indicated by: *^a^*, baseline and 24-hour time points in the air group; and *^b^*, baseline and 24-hour time points in the chlorine group.

BUN = blood urea nitrogen; ALT (GPT) = alanine aminotransferase; ALP = alkaline phosphatase; GGT = gamma-glutamyl transferase.

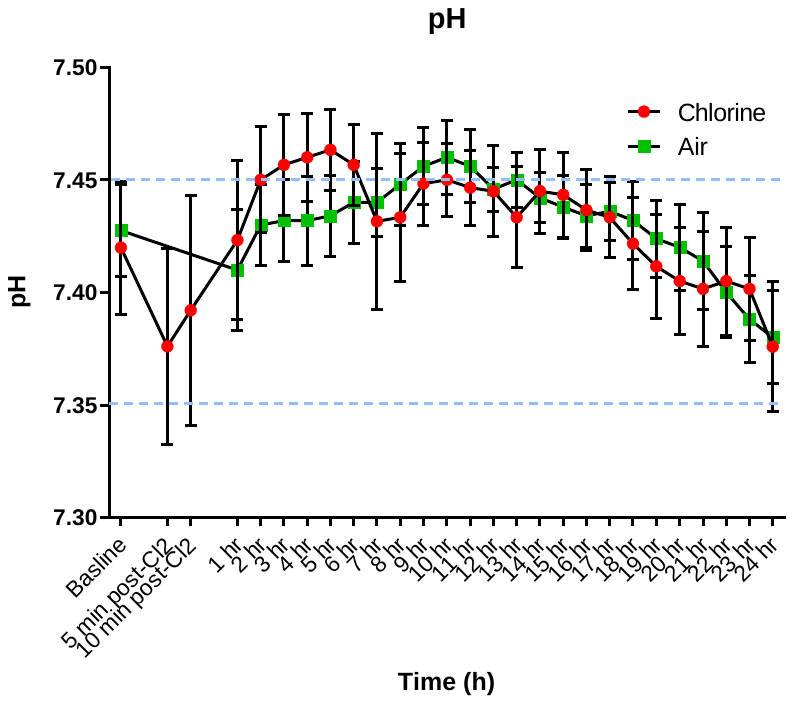
**Figure S1**

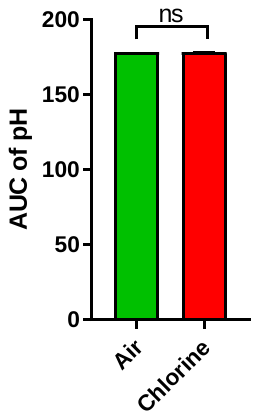

**Figure S1**: **Acid-base disturbance in chlorine-exposed pigs**. Anesthetized and mechanically ventilated pigs were exposed to either chlorine gas at ≤ 240 ppm or filtered room air for 1 hour. Arterial blood gas analysis was performed at hourly intervals. Figures S1A-D show changes in pH, the partial pressure of carbon dioxide (PaCO_2_), bicarbonate levels (HCO_3_^-^), and lactate levels in arterial blood over 24 hours, respectively. The blue dotted lines indicate range of respective normal values in humans.

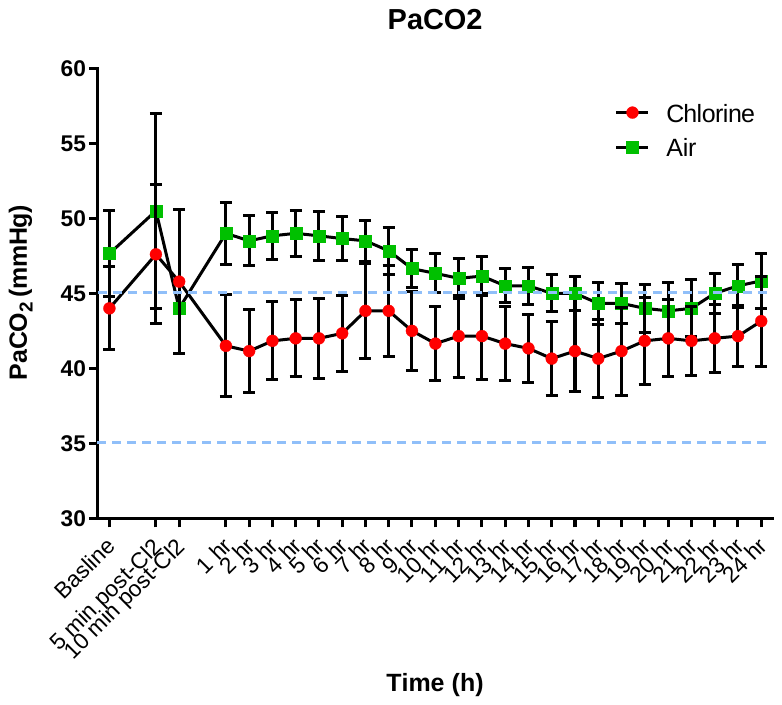

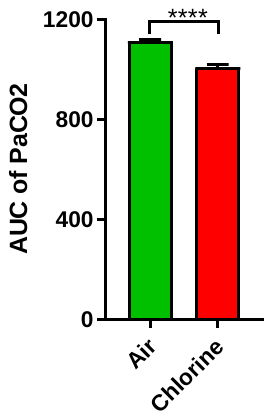

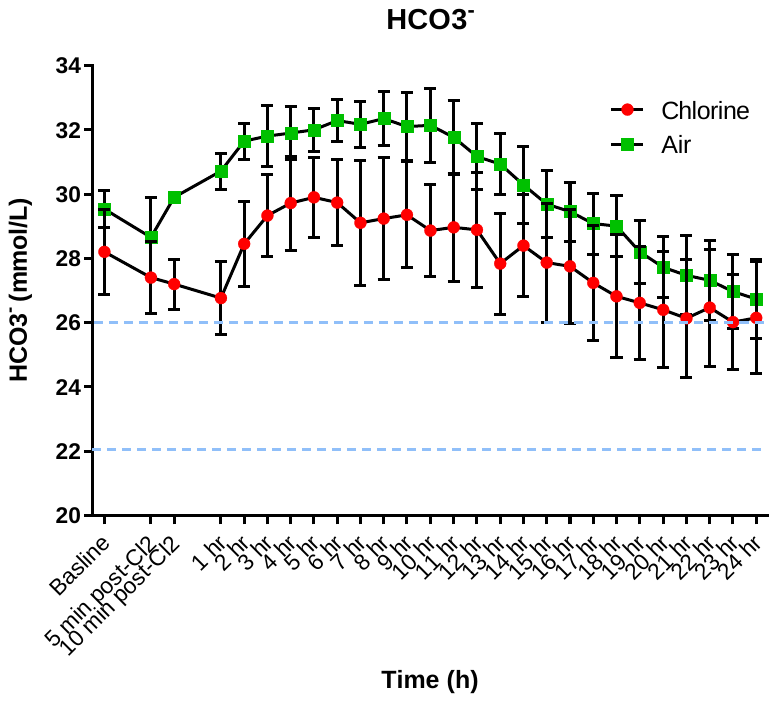

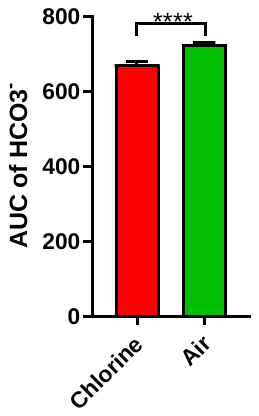

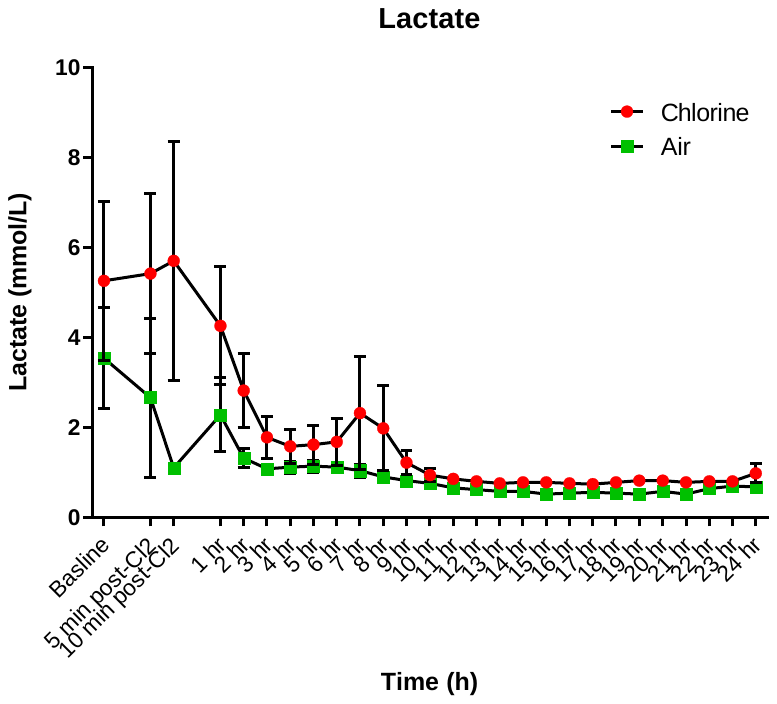

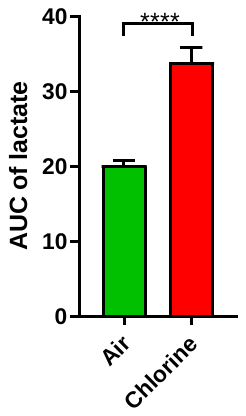

**Figure S2**

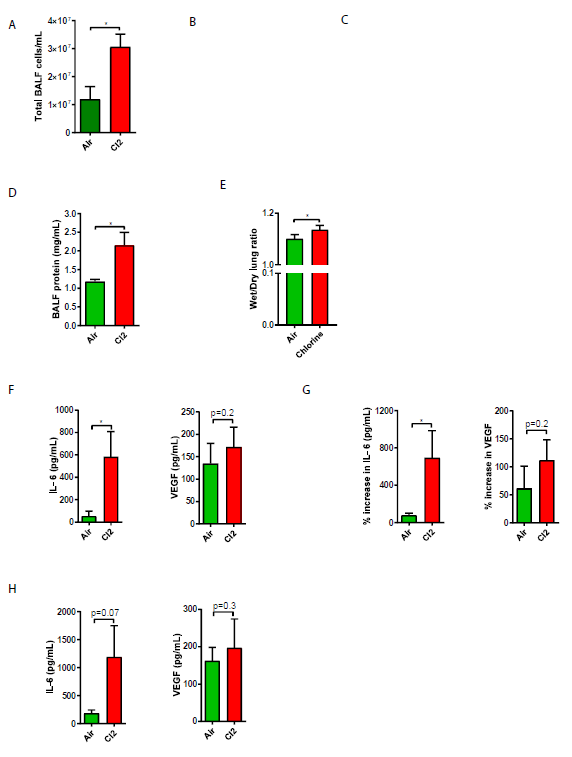

F

E

D

C B

B

**Figure S2**: **BALF cell counts, pulmonary vascular extravasation, edema, and pro-inflammatory cytokines in chlorine-exposed pigs.** Total BALF cell counts (Fig S2A), BALF protein (Fig S2B), wet/dry lung weight ratio (Fig S2C), pro-inflammatory cytokines (IL-6 and VEGF) in different biological matrices, ie, BALF (Fig S2D), plasma (Fig S2E), and lung tissue homogenate (Fig S2F) in Cl_2_-exposed or air-exposed pigs are presented. Unpaired *t*-tests were performed between chlorine- and air-exposed groups. Two-tailed *t*-tests were performed except for total BALF cell counts. Mann-Whitney test was performed to determine the significant difference in wet/dry weight lung ratio. *, *p* ≤ 0.05.

**Figure S3**

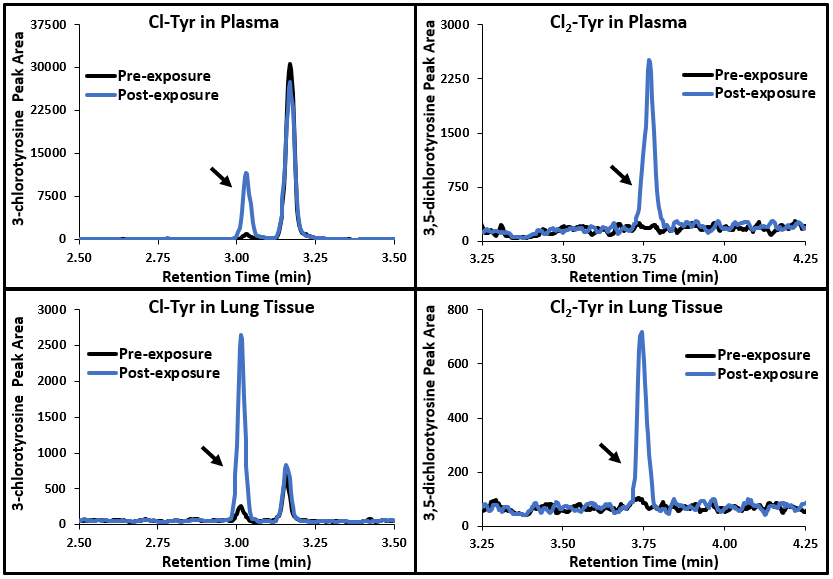

Figure S3 shows overlay of extracted ion chromatograms for plasma and lung samples collected from pigs pre- and post-chlorine gas exposure. The baseline (pre-exposure) is represented with a black line. Chlorine-exposed samples are indicated with a blue line. Arrows emphasize the Cl-Tyr and Cl_2_-Tyr peaks which have retention times at 3.1 min and 3.75 min, respectively.
